# Treatment-resistant depression and peripheral C-reactive protein

**DOI:** 10.1101/197012

**Authors:** Samuel R. Chamberlain, Jonathan Cavanagh, Peter de Boer, Valeria Mondelli, Declan Jones, Wayne C. Drevets, Philip J. Cowen, Neil A. Harrison, Linda Pointon, NIMA Consortium, Carmine M. Pariante, Edward T. Bullmore

**Affiliations:** Department of Psychiatry, School of Clinical Medicine, University of Cambridge, CB2 0SZ, UK; Cambridgeshire and Peterborough NHS Foundation Trust, Cambridge, CB21 5EF, UK; Sackler Centre, Institute of Health & Wellbeing, University of Glasgow, Sir Graeme Davies Building, Glasgow, G12 8TA, UK; Neuroscience, Janssen Research & Development, Janssen Pharmaceutica NV, Turnhoutseweg 30, B-2340, Beerse, Belgium; The Maurice Wohl Clinical Neuroscience Institute, Cutcombe Road, London, SE5 9RT, UK; Neuroscience, Janssen Research & Development, LLC, Titusville, NJ, 08560, USA; University Department of Psychiatry, Warneford Hospital, Oxford, OX3 7JX, UK; Brighton & Sussex Medical School, University of Sussex, Brighton, BN1 9RR, UK; Sussex Partnership NHS Foundation Trust, Swandean, BN13 3EP, UK; Stress, Psychiatry and Immunology Lab & Perinatal Psychiatry, Maurice Wohl Clinical Neuroscience Institute, Kings College, London, SE5 9RT, UK; Immuno-Psychiatry, Immuno-Inflammation Therapeutic Area Unit, GlaxoSmithKline R&D, Stevenage SG1 2NY, UK

## Abstract

**Research in context:** *Evidence before this study:* Dysregulation of the peripheral innate immune system has been implicated in the pathophysiology of major depressive disorder (MDD), and may partly account for why many patients do not experience symptomatic improvement. Elevated CRP has been demonstrated in meta-analysis for MDD compared to healthy volunteers, but little is known about whether this is the case for particular clinical phenotypes of the disorder, as opposed to MDD in general.

*Added value of this study:* This study recruited a large cohort of MDD patients, stratified by prior exposure to monoamine reuptake inhibitor treatment. MDD participants were carefully screened for physical comorbidity, and were compared to healthy volunteers matched for age, sex, body mass indices, and cigarette smoking status. Using group-wise comparisons and the innovative statistical approach of partial least squares, we demonstrated that elevated CRP was associated with treatment-resistance, childhood adversity, and specific depressive and anxious symptoms.

*Implications of all the available evidence:* CRP is significantly increased “on average” in MDD patients, However, CRP was most abnormally increased in the subgroup of patients with treatment-resistant depression. High BMI, high scores on vegetative symptoms of depression, low scores on calmness, and a history of childhood adversity, were all predictive of increased CRP. In future, stratification of MDD patients using pro-inflammatory biomarkers, like CRP, may be valuable for sample enrichment and targeted treatment interventions.

**Abstract:** *Background:* C-reactive protein (CRP) is a candidate biomarker for major depressive disorder (MDD), but it is unclear how peripheral CRP levels relate to the heterogeneous clinical phenotypes of the disorder.

*Methods:* We recruited 102 treatment-resistant, depressed MDD patients, 48 treatment-responsive, non-depressed MDD patients, 48 depressed but un-medicated patients, and 54 healthy volunteers. High sensitivity CRP in peripheral venous blood, body mass index (BMI), and questionnaire assessments of depression, anxiety, and childhood trauma, were measured. Group differences in CRP were estimated, before and after correction for BMI. Partial least squares (PLS) analysis explored the relationships between CRP and specific clinical phenotypes.

*Outcomes:* Compared to healthy volunteers, BMI-corrected CRP was significantly elevated in treatment-resistant patients (*P* = 0.007; Cohen’s d = 0.47); but not significantly so in the treatment-responsive (d = 0.29) and untreated (d = 0.18) groups. PLS yielded an optimal two factor solution that accounted for 34.7% of variation in clinical measures, and for 36.0% of variation in CRP. The clinical phenotypes most strongly associated with CRP and heavily weighted on the first PLS component were: vegetative depressive symptoms, BMI, state anxiety, and feeling unloved as a child or wishing for a different childhood.

*Interpretation:* Peripheral CRP was elevated in MDD, especially in treatment-resistant cases. Other phenotypes associated with elevated CRP included childhood adversity, and specific depressive and anxious symptoms. We suggest that MDD patients stratified for pro-inflammatory biomarkers, like CRP, have a distinctive clinical profile that might be responsive to second-line treatment with anti-inflammatory drugs.

*Funding:* Wellcome Trust strategy award to the Neuroimmunology of Mood Disorders and Alzheimer’s Disease (NIMA) Consortium.

## Introduction

Immunological mechanisms are increasingly implicated in the pathogenesis of depressive symptoms^1, 2^. Activation of the peripheral immune system has been consistently associated with major depressive disorder^3^. However, it has also been anticipated that not all patients with MDD will be peripherally inflamed to the same extent. A deeper understanding of how peripheral immune biomarkers relate to some of the dimensions of clinical heterogeneity encompassed by a diagnosis of MDD could be an important step towards mechanistically stratified treatment of depression in the future^2, 4^.

C-reactive protein (CRP) is an acute phase protein that is widely used in clinical practice and has also been measured in many prior studies of MDD. A high sensitivity assay for CRP is well validated and accessible. CRP synthesis is induced in the liver by pro-inflammatory cytokines – especially interleukin 6 (IL-6) – in response to infection, inflammation and tissue damage. In a meta-analysis of 20 case-control studies^3^, CRP was moderately increased “on average” (Cohen’s d = 0.47) in patients with MDD. However, there was significant heterogeneity of effect size between studies that may be attributable to clinical heterogeneity, with greater CRP in severe depression (Cohen’s d = 0.50) than in mild/moderate depression (Cohen’s d = 0.37), as well as methodological differences between studies^5^.

We were motivated to test the hypothesis that the clinically defined subgroup of patients with treatment-resistant depression would have the most abnormally increased CRP. An association between treatment resistance to monoaminergic anti-depressant drugs and increased CRP is hypothetically predictable on clinical and mechanistic grounds. Clinical studies indicate that pro-inflammatory cytokines that induce CRP synthesis are increased in treatment-resistant MDD. Pro-inflammatory cytokines can reduce the extracellular availability of serotonin by biasing expression of genes related to serotonin transport and tryptophan metabolism^6, 7^. Single studies have also reported that elevated CRP may be associated with other dimensions of clinical heterogeneity, *viz* atypical depression, childhood adversity, higher numbers of previous depressive episodes, or anxiety in male patients^2^.

We measured CRP in four groups of participants: currently depressed but not medicated (untreated) MDD patients; currently depressed and medicated (treatment-resistant) patients; currently medicated but not depressed (treatment-responsive) patients; and healthy volunteers with no history of MDD or monoaminergic drug treatment. The primary hypothesis, that CRP would be most clearly increased above normal levels in treatment-resistant patients with MDD, was tested by planned analyses of between-group differences in mean CRP. In a secondary analysis, we took a more exploratory approach to the question of what other dimensions of clinical heterogeneity in the sample might be related to variation in CRP. We used the multivariate technique of partial least squares (PLS) to explore the relationships between CRP and multiple (139) clinical phenotypes – ranging from BMI to questionnaire items for depressive symptoms, anxiety states or history of childhood adversity^8, 9^. In this way, we could identify a subset of clinical phenotypes weighted strongly on latent dimensions of clinical heterogeneity that were predictive of higher CRP levels. We also tested the confirmatory hypothesis that scores on these clinical dimensions of peripheral inflammation would be higher in the subgroup of patients with treatment resistance defined *a priori*.

### Patients and Methods

This was a non-interventional study, conducted as part of the Wellcome Trust Consortium for Neuroimmunology of Mood Disorders and Alzheimer’s disease (NIMA). There were five clinical study centres in the UK: Brighton, Cambridge, Glasgow, King’s College London, and Oxford. All procedures were approved by an independent Research Ethics Committee (National Research Ethics Service East of England, Cambridge Central, UK) and the study was conducted according to the Declaration of Helsinki. All participants provided informed consent in writing, and received £100 compensation for taking part.

#### Sample and eligibility criteria

We recruited four groups of participants: treatment-resistant depression, treatment-responsive depression, untreated depression, and healthy volunteers.

For all participants, the following inclusion criteria applied: age 25-50 years, able to give informed consent; able to fast for 8 hours, and abstain from strenuous exercise for 72h, prior to venous blood sampling; and fluent English. The following exclusion criteria applied: pregnancy or breast feeding; alcohol or substance use disorder in the preceding 12 months; participation in an investigational drug study within the preceding 12 months; lifetime history of any medical disorder or current use of any medication (e.g. statins, corticosteroids, anti-histamines, anti-inflammatory medications) likely to compromise interpretation of CRP (see **Supplemental Information**).

Adult patients meeting Diagnostic and Statistical Manual Version 5 (DSM-5) criteria for MDD were recruited from NHS mental health and primary care services and from the general population by purposive advertising. Lifetime histories of bipolar disorder or non-affective psychosis were additional exclusion criteria. Diagnosis of MDD and other psychiatric disorders was ascertained by Structured Clinical Interview. Current depressive symptom severity was defined using total scores from the 17-item Hamilton Rating Scale for Depression (HAM-D), and lifetime anti-depressant medication use was quantified using the Antidepressant Treatment Response Questionnaire (ATRQ). Citations for instruments are in **Supplemental Information.**

Patients were assigned to one of three subgroups or strata, per protocol: (i) treatment-resistant (DEP + MED +) patients who had total HAM-D > 13 and had been medicated with a monoaminergic drug at a therapeutic dose for at least six weeks; (ii) treatment-responsive (DEP-MED +) patients who had total HAM-D < 7 and had been medicated with a monoaminergic drug at a therapeutic dose for at least six weeks; and (iii) untreated (DEP + MED-) patients who had HAM-D > 17 and had not been medicated with a monoaminergic drug for at least six weeks. Cut-offs were defined *a priori* based on the literature. Total HAM-D score > 17 is a standard threshold for entry into placebo-controlled treatment trials of MDD; whereas a lower threshold of total HAM-D > 13 is typically used to define treatment-resistant depression, because there is usually some modest symptomatic response to treatment even if patients remain depressed^10, 11^.

A group of healthy volunteers was recruited by advertising with no current or past history of any major psychiatric disorder as defined by DSM-5, and no history of monoaminergic drug treatment for any indication. Healthy volunteers completed the same screening and baseline assessments as patient groups (see below).

Age, gender, medical history, smoking status, and family history were documented by semi-structured clinical interviews. Height and weight were measured for calculation of BMI (kg/m^2^).

#### Questionnaire assessments

Psychological symptoms and childhood adversity were assessed by administration of the following questionnaires (see **Supplemental Information**): the Beck depression inventory; the Spielberger State-Trait Anxiety Rating scale; the Chalder Fatigue Score; the Snaith-Hamilton Pleasure Scale; and the Childhood Trauma Questionnaire.

#### High sensitivity CRP measurement

High sensitivity C-reactive protein was measured as one of many immunological markers in a venous blood sample drawn from each participant. Here, we focus on CRP since this is convenient, and has been widely used^2^. Participants fasted for 8h, and abstained from strenuous exercise for 72h, prior to venous blood sampling between 08:00-10:00am.

#### Statistical analysis

For analysis of between-group differences in hs-CRP and other variables we first compared all MDD participants to healthy volunteers using planned t-tests. We then evaluated pairwise group differences using post-hoc t-tests, provided the main effect of group was significant by one-way analysis of variance (ANOVA). When assumptions of normality were violated, transforms and/or non-parametric tests were used. Cohen’s d was reported for the effect size of hs-CRP corrected for BMI in each clinical group compared to healthy volunteers. Additionally, we compared the proportion of participants in each group who had clinically elevated CRP, defined as > 3mg/L^12, 13^. The threshold for statistical significance was defined as two-tailed *P* < 0.05 throughout.

To identify demographic and clinical phenotypes associated with variation in CRP across all study participants, we utilized the method of partial least squares (PLS), as implemented in JMP Pro software Version 13.0^14^. PLS is a multivariate technique for modelling relationships between a set of predictor (*X*) and response (*Y*) variables in terms of a set of mutually orthogonal latent factors, or PLS components^9^. Detailed methodology is provided in **Supplemental Information**. The statistical significance of the final model was confirmed by comparing the percentage of variation in *X* and *Y* accounted for in the experimental data compared to the null distributions of the percentage of *X* or *Y* variance sampled by bootstrapping (1000 iterations).

## Results

### Demographic and clinical data

The size of each group and their demographic and clinical characteristics are summarized in **Table 1**. The groups did not differ significantly in terms of demographic characteristics. As expected, post hoc tests indicated that each group differed significantly from each other group on HAM-D total score (least significant t = 4.19, df = 248, P < 0.001). The mean number of failed pharmacological treatments for MDD episodes (<75% symptomatic response, defined by ATRQ) is listed for each clinical group in **Table 1**. The treatment-resistant group had more failed treatments than the untreated group (Wilcoxon Z = 2.843, P = 0.005); both the treatment-resistant group and the untreated group had significantly more failed treatments than the treatment-responsive group (Wilcoxon Z = 5.794, P < 0.001 and Wilcoxon Z = 3.079, P = 0.002, respectively). The majority of treatment-resistant patients were taking selective serotonin reuptake inhibitors (see **Table 1** footnote). Summary statistics for questionnaire-based measures and comorbidities are provided in **Tables S1** and **S2.**

**Table 1.**
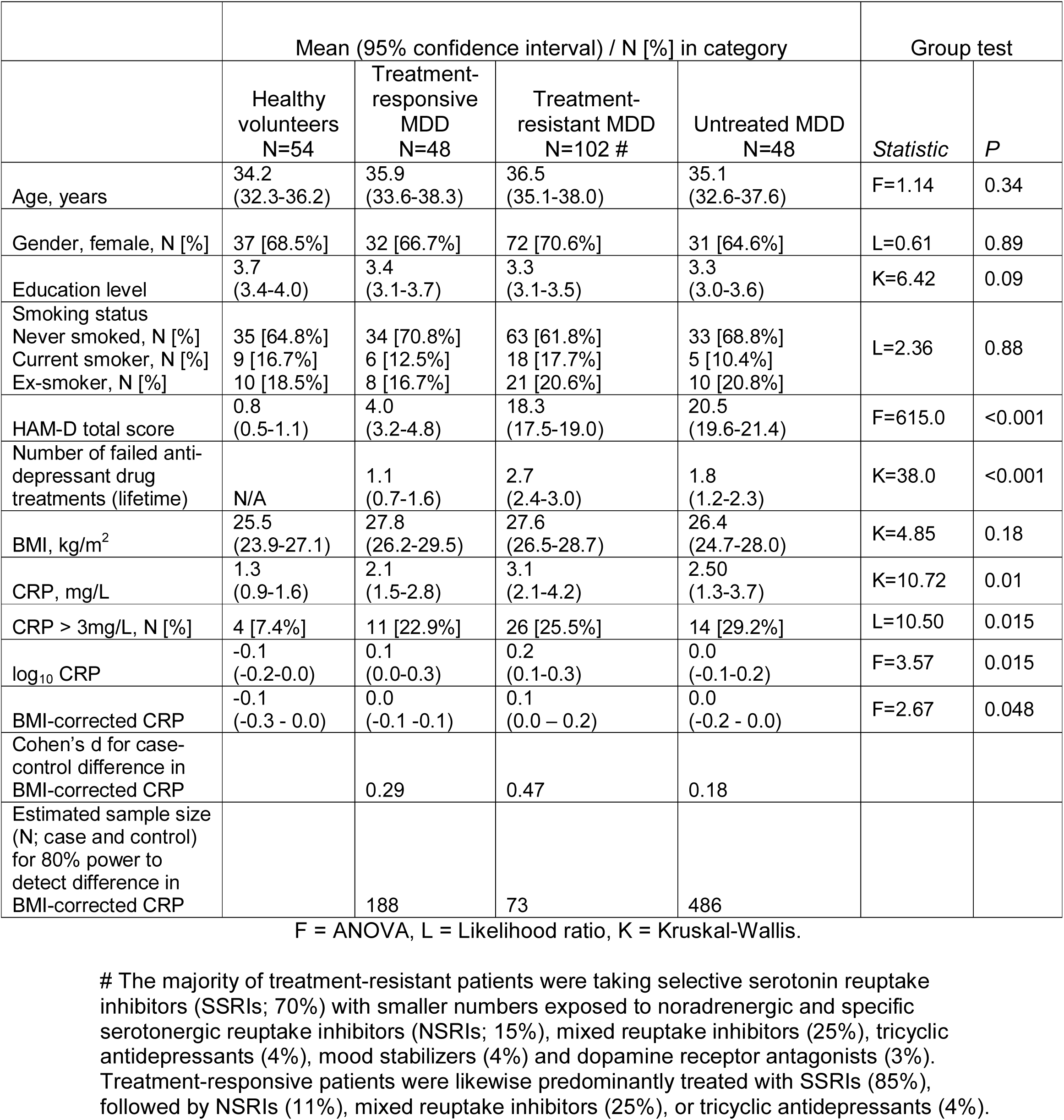
Demographic, clinical and hs-CRP data

|  | Mean (95% confidence interval) / N [%] in category |  |  |  | Group test |  |
| --- | --- | --- | --- | --- | --- | --- |
|  | Healthy volunteers<br>N=54 | Treatment-responsive<br>MDD<br>N=48 | Treatment-resistant MDD<br>N=102 # | Untreated MDD<br>N=48 | Statistic | P |
| Age, years | 34.2<br>(32.3-36.2) | 35.9<br>(33.6-38.3) | 36.5<br>(35.1-38.0) | 35.1<br>(32.6-37.6) | F=1.14 | 0.34 |
| Gender, female, N [%] | 37 [68.5%] | 32 [66.7%] | 72 [70.6%] | 31 [64.6%] | L=0.61 | 0.89 |
| Education level | 3.7<br>(3.4-4.0) | 3.4<br>(3.1-3.7) | 3.3<br>(3.1-3.5) | 3.3<br>(3.0-3.6) | K=6.42 | 0.09 |
| Smoking status |  |  |  |  |  |  |
| Never smoked, N [%] | 35 [64.8%] | 34 [70.8%] | 63 [61.8%] | 33 [68.8%] | L=2.36 | 0.88 |
| Current smoker, N [%] | 9 [16.7%] | 6 [12.5%] | 18 [17.7%] | 5 [10.4%] |  |  |
| Ex-smoker, N [%] | 10 [18.5%] | 8 [16.7%] | 21 [20.6%] | 10 [20.8%] |  |  |
| HAM-D total score | 0.8<br>(0.5-1.1) | 4.0<br>(3.2-4.8) | 18.3<br>(17.5-19.0) | 20.5<br>(19.6-21.4) | F=615.0 | <0.001 |
| Number of failed anti-depressant drug treatments (lifetime) | N/A | 1.1<br>(0.7-1.6) | 2.7<br>(2.4-3.0) | 1.8<br>(1.2-2.3) | K=38.0 | <0.001 |
| BMI, kg/m<sup>2</sup> | 25.5<br>(23.9-27.1) | 27.8<br>(26.2-29.5) | 27.6<br>(26.5-28.7) | 26.4<br>(24.7-28.0) | K=4.85 | 0.18 |
| CRP, mg/L | 1.3<br>(0.9-1.6) | 2.1<br>(1.5-2.8) | 3.1<br>(2.1-4.2) | 2.50<br>(1.3-3.7) | K=10.72 | 0.01 |
| CRP > 3mg/L, N [%] | 4 [7.4%] | 11 [22.9%] | 26 [25.5%] | 14 [29.2%] | L=10.50 | 0.015 |
| log<sub>10</sub> CRP | -0.1<br>(-0.2-0.0) | 0.1<br>(0.0-0.3) | 0.2<br>(0.1-0.3) | 0.0<br>(-0.1-0.2) | F=3.57 | 0.015 |
| BMI-corrected CRP | -0.1<br>(-0.3 - 0.0) | 0.0<br>(-0.1 -0.1) | 0.1<br>(0.0 – 0.2) | 0.0<br>(-0.2 - 0.0) | F=2.67 | 0.048 |
| Cohen's d for case-control difference in BMI-corrected CRP |  | 0.29 | 0.47 | 0.18 |  |  |
| Estimated sample size (N; case and control) for 80% power to detect difference in BMI-corrected CRP |  | 188 | 73 | 486 |  |  |

F = ANOVA, L = Likelihood ratio, K = Kruskal-Wallis.

#### C-reactive protein

Mean hs-CRP concentrations (and 95% confidence intervals) are shown in **Table 1**. Mean CRP was significantly increased in all MDD cases compared to healthy controls (Wilcoxon Z = 2.7, P = 0.007). Both treatment-resistant and treatment-responsive groups had significantly higher mean hs-CRP than controls (Wilcoxon *Z* = 2.9, *P* = 0.004 and Wilcoxon *Z* = 2.6, *P* = 0.010, respectively).

The proportion of participants with hs-CRP levels exceeding the conventional threshold value of 3mg/L was also significantly different between the pooled MDD groups and controls (likelihood ratio chi-square (LR) = 10.01, P = 0.002). Treatment-resistant, untreated, and treatment-responsive MDD groups all had significantly increased proportions of participants with hs-CRP > 3 mg/L compared to healthy volunteers (LR = 8.4, *P* = 0.004; LR = 8.6, *P* = 0.003; and LR = 5.0, *P* = 0.025, respectively). No other post hoc test was statistically significant; i.e. depressed groups did not differ significantly from each other (all *P* > 0.09).

#### Log-transformed and BMI-corrected C-reactive protein

The distributions of hs-CRP were positively skewed (moment skewness = 5.08) and therefore were normalized by base 10 log transform; see **Figure 1**. Log_10_ CRP was significantly increased in all MDD cases compared to controls (t = 2.81, df = 250, P = 0.004). Only the treatment-resistant and treatment-responsive cases had significantly higher log_10_ CRP than controls (*t* = 3.07, df = 248, *P* = 0.002 and *t* = 2.32, df = 248, *P* = 0.021, respectively).

**Figure 1.**
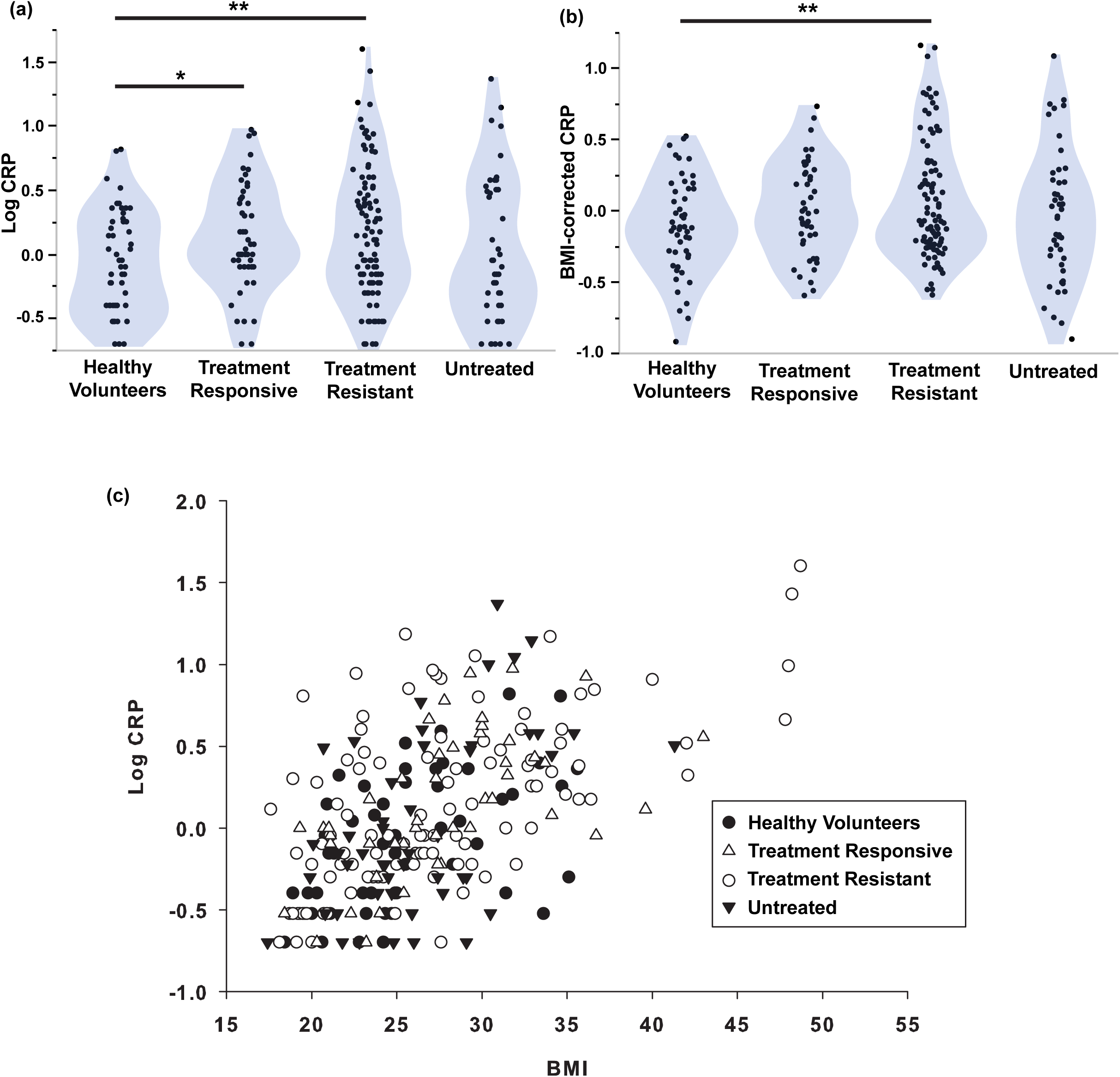
High sensitivity CRP and its relationship with BMI. **(a)**: Violin plots of log_10_ CRP for each of three subgroups of MDD patients and healthy volunteers. **(b)**: Violin plots of BMI-corrected CRP (log_10_ CRP regressed on log_10_ BMI) for each of three subgroups of MDD patients and healthy volunteers. **(c)**: Scatterplot of BMI versus log_10_ CRP (Spearman’s rho = + 0.57, P < 0.001), with points coded by sample group. The main effects of group were significant for (a) and (b) (ANOVA, *F* = 3.57, *P* = 0.015; *F* = 2.67, *P* = 0.048). * *P* < 0.05, ** *P* < 0.01 significant pair-wise difference between groups by post hoc *t*-tests.

As anticipated by prior studies, there was a significant positive correlation between BMI and log_10_ CRP across all study participants (Spearman’s rho = 0.56, df = 250, *P* < 0.001; **Figure 1**). Since BMI data were also positively skewed (moment skewness = 1.03) ^15^, we regressed log_10_ CRP on log_10_ BMI and used the residuals as estimates of BMI-corrected CRP (**Figure 1**). BMI-corrected CRP was significantly elevated in all MDD cases compared to controls (t = 2.24, df = 238, P = 0.026). Post hoc *t*-tests indicated that only the treatment-resistant cases had significantly higher mean BMI-corrected CRP than the controls (*t* = 2.71, df = 236, *P* = 0.007; Cohen’s d = 0.47).

To assess the possible confounding effect of symptom severity, we identified the subgroup of treatment-resistant patients (N=48) that had total HAM-D > 17, thereby corresponding to the cut-off used to define the untreated group. We confirmed that BMI-corrected CRP was abnormally increased in treatment-resistant patients with HAM-D > 18 (t = 3.0, P = 0.004) with a case-control difference of similar size (Cohen’s d = 0.43) to that of treatment-resistant patients with HAM-D >13.

#### Partial least squares analysis of the relationship between CRP and clinical variables

Thirteen (out of 139) clinical phenotypes passed criterion for an important effect on CRP levels. Iterative cross-validation of the PLS model including only these important variables yielded an optimal two-factor solution (**Figure 2 and S2**), which accounted in total for 34.7% of variation in clinical measures (*X*), and for 36.1% of variation in CRP (*Y*). This differed significantly from the proportions of variance expected under the null hypothesis (% Var (*X*) = 12.3%, 95% CI 12.1-12.5%; % Var (*Y*) = 2.7%, 95% CI 1.9-3.0%).

**Figure 2.**
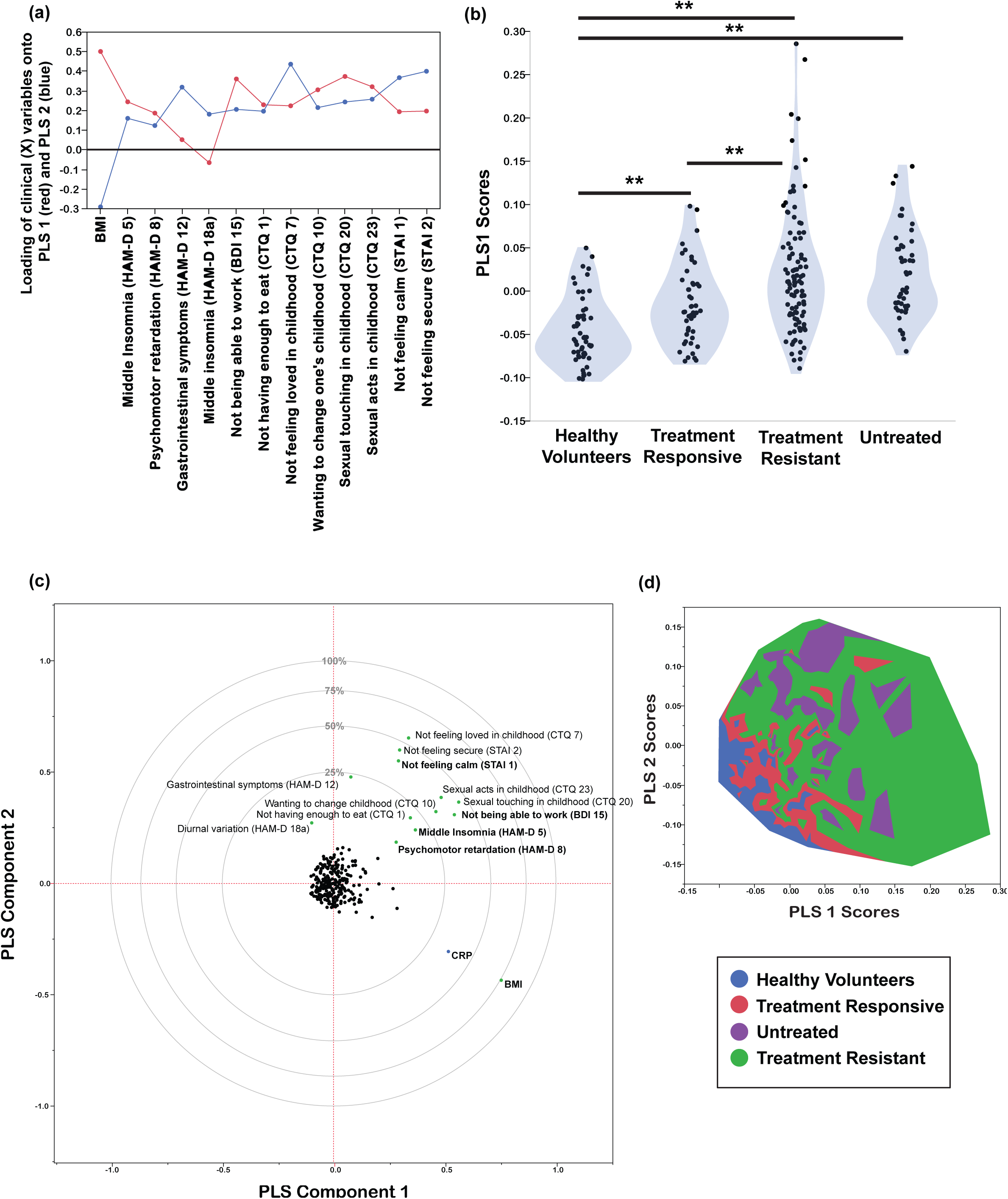
Partial least squares analysis of the relationships between hs-CRP and clinical phenotypes. **(a)**: Loadings of clinical (*X*) variables onto the first two PLS components. These 13 variables passed the first criterion for an important effect on CRP variance. **(b)**: Violin plot of PLS1 X scores for each of three subgroups of MDD patients and healthy volunteers. ** *P* < 0.01 significant pair-wise difference between groups by post hoc *t -* tests. **(c)**: Plot of the clinical (*X*) and CRP (*Y*) variables in the space of the first two PLS components. The clinical variables named in bold font passed both criteria for an important effect on CRP; the variables named in normal font passed the first criterion but not the more conservative second criterion. Panel (d): Contour plot for distribution of study participants in the space of the first two PLS components, color-coded by group, confirming that patients with treatment-resistant depression had high scores on the clinical syndrome of variables represented by PLS1.

The first PLS component (PLS1) accounted for 26.7% of the variation in hs-CRP. Positive scores on PLS1 indicated higher CRP. The clinical phenotypes that were significantly weighted on PLS1 were higher BMI, not feeling loved in childhood (CTQ item 7), not feeling calm (STAI item 1), wanting to change one’s family in childhood (CTQ item 10), psychomotor retardation (HAM-D item 8), middle insomnia (HAM-D item 5) and not being able to work (BDI item 15). The PLS1 scores for individual participants differed significantly between groups (*F* = 19.88, df = 3,248, *P* < 0.001; **Figure 2**). PLS1 scores were highest in the treatment-resistant group, followed by the untreated group, the treatment-responsive group and the healthy volunteers.

The second PLS component (PLS2) orthogonally explained 9.4% of variation in CRP. Positive scores on PLS 2 indicated lower CRP. PLS2 scores differed significantly between groups (*F* = 24.34, df = 3,248, *P* < 0.001; **Figure S3**). Untreated patients had the highest PLS2 scores, followed by treatment-resistant patients, treatment-responsive patients and healthy volunteers.

## Discussion

This is the first study to measure peripheral C-reactive protein using the same high sensitivity assay across a large sample of MDD patients (N=198) prospectively stratified in terms of their current and past history of treatment with monoaminergic antidepressant drugs. We replicated the well-established finding that CRP is significantly increased “on average” in MDD patients, screened for physical comorbidity, and compared to healthy volunteers matched for age, sex, BMI and cigarette smoking status. However, we also found evidence for our primary hypothesis that CRP was most abnormally increased in the subgroup of patients with treatment-resistant depression (N=102). The standardized size of the case-control difference in CRP between healthy volunteers and treatment-resistant cases (Cohen’s d = 0.47) was greater than the case-control difference for treatment-responsive cases (0.29), or untreated cases (0.18). Controlling for non-normality of the CRP distribution, and for the strong positive correlation between CRP and obesity, we found that the case-control difference in CRP remained significant only for the subgroup of treatment-resistant cases. These results of planned analysis are consistent with the hypothesis that peripheral inflammation is a marker or risk factor for treatment-resistant depression.

Taking a convergent but more exploratory approach to the data, we used multivariate analysis to identify two dimensions of clinical heterogeneity that were predictive of CRP. We found that a subset of 15 out of 131 clinically measured phenotypes explained ∽ 36% of the variance in CRP. High BMI, high scores on vegetative symptoms of depression, low scores on calmness, and a history of childhood adversity, were all predictive of increased CRP. As expected from the results of our primary analysis, we confirmed that the group of patients defined *a priori* in terms of treatment resistance had the highest scores on this clinical profile associated with high CRP.

### Treatment-resistant depression and peripheral inflammation

Monoamine reuptake inhibitors and related drugs are evidence-based pharmacological treatments for MDD; but response failure afflicts approximately 30% of patients^16^. Because of the global burden of disability attributable to MDD, the search for improved understanding of biomarkers of therapeutic resistance to current first-line treatment options is pressing. Our results provide fresh evidence that patients with treatment-resistant depression have the most abnormally increased CRP levels compared both to treatment-responsive and currently untreated patients. To our knowledge, a specific relationship between CRP and monoaminergic antidepressant drug treatment resistance has not been demonstrated previously, although there is evidence both for increased pro-inflammatory cytokine concentrations^2^ and for increased peripheral expression of cytokine related genes^17^ in treatment-resistant depression. There is also some evidence that baseline inflammatory markers may be useful predictors of treatment response in MDD^18^. In a rat model of treatment-resistant depression, elevated CRP at baseline differentiated responders from non-responders to ketamine, an NMDA receptor antagonist with anti-inflammatory and anti-depressant effects^19^. At a cellular level, neurons, microglia and macrophages respond to inflammatory challenges by activating metabolic pathways that reduce the synaptic availability of serotonin and catalyse the conversion of tryptophan to kynurenine and its putatively neurotoxic, glutamatergic agonist metabolites^7^. These effects of inflammation on serotonin transport and tryptophan metabolism may constitute a mechanism by which peripheral inflammation is associated with lack of therapeutic response to SSRIs^20^.

### Clinical phenotypes predictive of increased CRP in depression

Obesity and its cardiovascular sequelae have been repeatedly associated with increased CRP. In the current study, which excluded patients with a lifetime history of medical disorders including atherosclerosis and diabetes, we confirmed that higher BMI was strongly associated with higher CRP levels. One mechanistic explanation is that macrophages constitute up to 60% of cells in adipose tissue and can release large amounts of IL-6, which is a key driver of CRP synthesis^21^. So it is not surprising that inflammation (CRP) and obesity (BMI) were related herein; however, we do not consider that this association trivially accounts for increased CRP in treatment-resistant depression. The groups did not differ significantly in baseline BMI and the case-control difference remained significant for the treatment-resistant patients even after statistical regression to control for individual differences in BMI.

Of all the depressive symptoms measured, so-called vegetative symptoms (psychomotor retardation, insomnia, difficulty getting started/difficulty working) were more important in explaining higher CRP. These findings are consistent with prior reports that somatic but not cognitive symptoms of depression were associated with increased CRP^22^. Vegetative symptoms of depression are akin to the illness or sickness behavior that has been repeatedly demonstrated in animal models and experimental medicine studies of humans exposed to acute pro-inflammatory challenge^23^. We also found evidence that state anxiety was related with CRP, which is compatible with prior data linking acute endotoxin exposure to anxious and depressive states in healthy volunteers^24^.

It is established that childhood trauma increases risk of later mental health disorders, including depression^25^. In a meta-analysis, individuals exposed to childhood trauma had significantly elevated levels of CRP in adulthood, albeit with a small effect size (Fisher’s Z = 0.10)^26^. In a longitudinal study of female adolescents at risk of depression, childhood adversity was found to promote subsequent clustering of depression and inflammation^27^. These results are compatible with our findings that feeling unloved in childhood and wanting to change one’s family in childhood were significantly correlated with higher CRP in adults.

### Methodological issues

Due to the case-control design, between-group differences in CRP could theoretically be confounded by other factors influencing peripheral inflammation. However, we excluded patients with inflammatory disorders or anti-inflammatory drug treatment; and the groups were matched for demographic characteristics. The lack of statistically significant case-control differences in BMI-corrected CRP for the comparisons between healthy volunteers and the treatment-responsive and untreated MDD groups could theoretically reflect the smaller sizes of these groups compared to the treatment-resistant MDD group. However, power calculations indicated that the case-control differences in BMI-corrected CRP would probably not have been significant even if the treatment-responsive group had the same size as the resistant group (**Table 1**). The depression symptom severity threshold or cut-off score for the treatment-resistant group was HAM-D > 13, whereas the cut-off score for the untreated group was HAM-D > 18^10, 11^. Potentially the resulting difference in symptom severity between these groups could explain the greater increase of CRP in the treatment-resistant group compared to controls. However, matching the cut-off used for the untreated group made little difference to the primary results. Treatment resistance was defined by inadequate response to the current drug treatment whereas some other criteria for treatment resistance stipulate failed response to at least two drugs of different mechanisms of action. The study was not planned or powered to test differences in CRP between subgroups defined by dose or type of current anti-depressant medication. The sample was recruited from the UK population, which is known to differ from the US and other populations in terms of BMI and other factors that can influence the numerical distribution of CRP and this may mitigate generalizability of our results. Finally, CRP is only one of many markers of peripheral inflammation that have been, or could be, linked to depression. Although these data demonstrate that CRP is robustly associated with treatment-resistant depression, we do not claim that CRP is necessarily the best of all possible peripheral blood biomarkers of treatment-resistant depression.

### Conclusions

Major depressive disorder is associated with increased CRP compared to healthy volunteers and the case-control difference is greatest in treatment-resistant depression. Increased CRP and treatment resistance were also associated with other aspects of clinical heterogeneity in depression including obesity, vegetative symptoms of fatigue and sleep disturbance, state anxiety, and a history of childhood adversity. We suggest there may be a clinically and immunologically diagnosable sub-syndrome of “inflamed depression” comprising the MDD patients most likely to benefit therapeutically from second-line treatment with anti-inflammatory drugs^13^.

## Funding and Disclosure

This work was funded by a Wellcome Trust strategy award to the Neuroimmunology of Mood Disorders and Alzheimer’s Disease (NIMA) Consortium which is also funded by Janssen, GlaxoSmithKline, Lundbeck and Pfizer. Recruitment of patients was supported by the National Institute of Health Research (NIHR) Clinical Research Network: Kent, Surrey and Sussex & Eastern. SRC consults for Cambridge Cognition and Shire; and his input in this project was funded by a Wellcome Trust Clinical Fellowship (110049/Z/15/Z). ETB is employed half-time by the University of Cambridge and half-time by GlaxoSmithKline; he holds stock in GSK. In the last three years PJC has served on an advisory board for Lundbeck. NAH consults for GSK. PdB, DJ and WCD are employees of Janssen Research & Development, LLC., of Johnson & Johnson, and hold stock in Johnson & Johnson. The other authors report no financial disclosures or potential conflicts of interest.

### Acknowledgements

The authors would like to thank all study participants, research teams, and laboratory staff, without whom this research would not have been possible.

Members of the NIMA Consortium are thanked and acknowledged:

## Cambridge

Edward T. Bullmore (PI, EC)^1,2,11^, Junaid Bhatti^1^, Samuel R. Chamberlain^1,2^, Marta M. Correia^1,12^, Amber Dickinson*, Andy Foster^2^, Manfred Kitzbichler^1^, Clare Knight^2^, Mary-Ellen Lynall^1^, Christina Maurice^1^, Howard Mount^13^, Ciara O’Donnell^1^, Linda J. Pointon^1^, Peter St George Hyslop^1,13,14^, Lorinda Turner^1^, Barry Widmer^1^, Guy B. Williams^1,14^

## Cardiff

B. Paul Morgan (PI)^15^, Claire Leckey^15^, Angharad Morgan^15^, Caroline O’Hagan*, Samuel Touchard^15^

## Glasgow

Jonathan Cavanagh (PI, EC)^3^, Catherine Deith*, John McClean^16^, Alison McColl^3^, Andrew McPherson*, Paul Scouller*, Murray Sutherland^16^

## Independent advisor

H.W.G.M. (Erik) Boddeke (EC)^17^

## GSK

Jill Richardson (EC)^18^, Shahid Khan^11^, Phil Murphy^19^, Christine Parker^19^, Jai Patel^11^

## Janssen

Declan Jones (EC)^6^, Peter de Boer^4^, John Kemp^4^, Paul Acton^6^, Wayne C. Drevets^6^, Jeffrey S. Nye (deceased), Gayle Wittenberg^6^, John Isaac^6^, Anindya Bhattacharya^6^, Nick Carruthers^6^, Hartmuth Kolb^6^

## Kings College London

Carmine Pariante (PI)^10^, Gareth Barker^20^, Heidi Byrom^10^, Diana Cash^20^, Antony Gee^20^, Caitlin Hastings^10^, Nicole Mariani^10^, Anna McLaughlin^10^, Valeria Mondelli^10^, Maria Nettis^10^, Naghmeh Nikkheslat^10^, Karen Randall^20^, Hannah Sheridan*, Camilla Simmons^20^, Nisha Singh^20^, Federico Turkheimer^20^, Victoria Van Loo*, Marta Vicente Rodriguez^20^, Tobias Wood^20^, Courtney Worrell*, Zuzanna Zajkowska*

## Lundbeck

Niels Plath (EC)^21^, Jan Egebjerg^21^, Hans Eriksson^21^, Francois Gastambide^21^, Karen Husted Adams^21^, Ross Jeggo^21^, Christian Thomsen^21^, Jason O’Connor^22^, Jan Torleif Pederson^21^, Brian Campbell*, Thomas Möller*, Bob Nelson*, Stevin Zorn*

## Oxford

Mary Jane Attenburrow (PI)^7,23^, Alison Baird, Jithen Benjamin^23^, Stuart Clare^25^, Philip Cowen^7^, I-Shu (Dante) Huang^24^, Samuel Hurley*, Helen Jones^23^, Simon Lovestone^7^, Francesca Mada^23^, Alejo Nevado-Holgado^7^, Akintayo Oladejo*, Elena Ribe^7^, Anviti Vyas*

## Pfizer

Zoe Hughes (EC)^26^, Rita Balice-Gordon*, Brendon Binneman^26^, James Duerr^26^, Terence Fullerton^26^, Justin Piro^26^, Tarek Samad^26^, Jonathan Sporn^26^

## Southampton

Hugh Perry (PI)^27^, Madeleine Cleal*, Gemma Fryatt^27^, Diego Gomez-Nicola^27^, Renzo Mancuso^27^

## Sussex

Neil Harrison (PI, EC)^28^, Mara Cercignani^28^, Charlotte Clarke^28^, Elizabeth Hoskins^29^, Charmaine Kohn^29^, Rosemary Murray*, Dominika Wlazly^30^

PI = Principal Investigator

EC = Executive committee member

## Footnotes

1 Department of Psychiatry, School of Clinical Medicine, University of Cambridge, CB2 0SZ, UK

2 Cambridgeshire and Peterborough NHS Foundation Trust, Cambridge, CB21 5EF, UK

3 Sackler Centre, Institute of Health & Wellbeing, University of Glasgow, Sir Graeme Davies Building, Glasgow, G12 8TA, UK

4 Neuroscience, Janssen Research & Development, Janssen Pharmaceutica NV, Turnhoutseweg 30, B-2340, Beerse, Belgium

5 The Maurice Wohl Clinical Neuroscience Institute, Cutcombe Road, London, SE5 9RT, UK

6 Neuroscience, Janssen Research & Development, LLC, Titusville, NJ, 08560, USA

7 University Department of Psychiatry, Warneford Hospital, Oxford, OX3 7JX, UK

8 Brighton & Sussex Medical School, University of Sussex, Brighton, BN1 9RR, UK

9 Sussex Partnership NHS Foundation Trust, Swandean, BN13 3EP, UK

10 Stress, Psychiatry and Immunology Lab & Perinatal Psychiatry, Maurice Wohl Clinical Neuroscience Institute, Kings College, London, SE5 9RT, UK

11 Immuno-Psychiatry, Immuno-Inflammation Therapeutic Area Unit, GlaxoSmithKline R&D, Stevenage SG1 2NY, UK

12 MRC Cognition and Brain Sciences Unit, 15 Chaucer Road, Cambridge CB2 7EF, UK

13 Tanz Centre for Research in Neurodegenerative Diseases, 60 Leonard Avenue, Toronto, ON M5T 2S8 Canada

14 Department of Clinical Neurosciences, University of Cambridge, CB2 0SZ, UK

15 University of Cardiff, Cardiff CF10 3AT, UK

16 NHS Greater Glasgow and Clyde, 1055 Great Western Rd, Glasgow G12 0XH, UK

17 University of Groningen, 9712 CP Groningen, Netherlands

18 Neurosciences Virtual PoC DPU, GlaxoSmithKline R&D, Stevenage SG1 2NY, UK

19 Experimental Medicine Imaging, GlaxoSmithKline R&D, Stevenage SG1 2NY, UK

20 Centre for Neuroimaging Sciences, Denmark Hill, London SE5 9AF, UK

21 H. Lundbeck A/S Ottiliavej 9, 2500, Valby, Denmark

22 University of Texas Health Science Center at San Antonio, 7703 Floyd Curl Dr, San Antonio, TX 78229, USA

23 NIHR Oxford cognitive health Clinical Research Facility, Warneford Hospital, Oxford, OX3 7JX, UK

24 The Kennedy Institute of Rheumatology, Roosevelt Dr, Oxford OX3 7FY, UK

25 Oxford Centre for Functional MRI of the Brain, John Radcliffe Hospital, Oxford OX3 9DU, UK

26 Pfizer, Inc, 1 Portland Street, Cambridge MA, USA

27 Centre for Biological Sciences, University of Southampton, Southampton, UK

28 Clinical Imaging Sciences Centre (CISC), University of Sussex, Brighton, BN1 9RR, UK

29 Sussex Partnership NHS Foundation Trust, Nevill Avenue, Hove BN3 7HZ, UK

30 Brighton & Sussex University Hospitals NHS Trust, Brighton BN2 5BE, UK

* Former consortium members

## References

1. Dantzer R, O’Connor JC, Freund GG, Johnson RW, Kelley KW. From inflammation to sickness and depression: when the immune system subjugates the brain. Nat Rev Neurosci. 2008;9(1):46–56.

2. Strawbridge R, Arnone D, Danese A, Papadopoulos A, Herane Vives A, Cleare AJ. Inflammation and clinical response to treatment in depression: A meta-analysis. Eur Neuropsychopharmacol. 2015;25(10):1532–43.

3. Haapakoski R, Mathieu J, Ebmeier KP, Alenius H, Kivimaki M. Cumulative meta-analysis of interleukins 6 and 1beta, tumour necrosis factor alpha and C-reactive protein in patients with major depressive disorder. Brain Behav Immun. 2015;49:206–15.

4. Domenici E, Wille DR, Tozzi F, Prokopenko I, Miller S, McKeown A, et al. Plasma protein biomarkers for depression and schizophrenia by multi analyte profiling of case-control collections. PLoS One. 2010;5(2):e9166.

5. Glaser R, Robles TF, Sheridan J, Malarkey WB, Kiecolt-Glaser JK. Mild depressive symptoms are associated with amplified and prolonged inflammatory responses after influenza virus vaccination in older adults. Arch Gen Psychiatry. 2003;60(10):1009–14.

6. Janssen DG, Caniato RN, Verster JC, Baune BT. A psychoneuroimmunological review on cytokines involved in antidepressant treatment response. Hum Psychopharmacol. 2010;25(3):201–15.

7. Leday GGR, Vertes PE, Richardson S, Greene JR, Regan T, Khan S, et al. Replicable and Coupled Changes in Innate and Adaptive Immune Gene Expression in Two Case-Control Studies of Blood Microarrays in Major Depressive Disorder. Biol Psychiatry. 2017.

8. Abdi H, Williams LJ. Partial least squares methods: partial least squares correlation and partial least square regression. Methods Mol Biol. 2013;930:549–79.

9. Wold H. Estmation of principal components and related models by iterative least squares. New York: Academic Press; 1966.

10. Marcus RN, McQuade RD, Carson WH, Hennicken D, Fava M, Simon JS, et al. The efficacy and safety of aripiprazole as adjunctive therapy in major depressive disorder: a second multicenter, randomized, double-blind, placebo-controlled study. J Clin Psychopharmacol. 2008;28(2):156–65.

11. Katona CL, Robertson MM, Abou-Saleh MT, Nairac BL, Edwards DR, Lock T, et al. Placebo-controlled trial of lithium augmentation of fluoxetine and lofepramine. Int Clin Psychopharmacol. 1993;8(4):323.

12. Bassuk SS, Rifai N, Ridker PM. High-sensitivity C-reactive protein: clinical importance. Curr Probl Cardiol. 2004;29(8):439–93.

13. Raison CL, Rutherford RE, Woolwine BJ, Shuo C, Schettler P, Drake DF, et al. A randomized controlled trial of the tumor necrosis factor antagonist infliximab for treatment-resistant depression: the role of baseline inflammatory biomarkers. JAMA Psychiatry. 2013;70(1):31–41.

14. JMP Pro ® Version 13.0, Cary, North Carolina, USA. SAS Institute Inc.; 2017.

15. Terpening WD. Statistical analysis for business using JMP: A Student’s Guide. Cary, North Carolina, USA: SAS Institute Inc.; 2011.

16. Al-Harbi KS. Treatment-resistant depression: therapeutic trends, challenges, and future directions. Patient Prefer Adherence. 2012;6:369–88.

17. Cattaneo A, Gennarelli M, Uher R, Breen G, Farmer A, Aitchison KJ, et al. Candidate genes expression profile associated with antidepressants response in the GENDEP study: differentiating between baseline ‘predictors’ and longitudinal ‘targets’. Neuropsychopharmacology. 2013;38(3):377–85.

18. Jha MK, Minhajuddin A, Gadad BS, Greer T, Grannemann B, Soyombo A, et al. Can C-reactive protein inform antidepressant medication selection in depressed outpatients? Findings from the CO-MED trial. Psychoneuroendocrinology. 2017;78:105–13.

19. Walker AJ, Foley BM, Sutor SL, McGillivray JA, Frye MA, Tye SJ. Peripheral proinflammatory markers associated with ketamine response in a preclinical model of antidepressant-resistance. Behav Brain Res. 2015;293:198–202.

20. Haroon E, Raison CL, Miller AH. Psychoneuroimmunology meets neuropsychopharmacology: translational implications of the impact of inflammation on behavior. Neuropsychopharmacology. 2012;37(1):137–62.

21. Mohamed-Ali V, Goodrick S, Rawesh A, Katz DR, Miles JM, Yudkin JS, et al. Subcutaneous adipose tissue releases interleukin-6, but not tumor necrosis factor-alpha, in vivo. J Clin Endocrinol Metab. 1997;82(12):4196–200.

22. Duivis HE, Vogelzangs N, Kupper N, de Jonge P, Penninx BW. Differential association of somatic and cognitive symptoms of depression and anxiety with inflammation: findings from the Netherlands Study of Depression and Anxiety (NESDA). Psychoneuroendocrinology. 2013;38(9):1573–85.

23. Capuron L, Gumnick JF, Musselman DL, Lawson DH, Reemsnyder A, Nemeroff CB, et al. Neurobehavioral effects of interferon-alpha in cancer patients: phenomenology and paroxetine responsiveness of symptom dimensions. Neuropsychopharmacology. 2002;26(5):643–52.

24. Reichenberg A, Yirmiya R, Schuld A, Kraus T, Haack M, Morag A, et al. Cytokine-associated emotional and cognitive disturbances in humans. Arch Gen Psychiatry. 2001;58(5):445–52.

25. Sachs-Ericsson N, Kendall-Tackett K, Hernandez A. Childhood abuse, chronic pain, and depression in the National Comorbidity Survey. Child Abuse Negl. 2007;31(5):531–47.

26. Baumeister D, Akhtar R, Ciufolini S, Pariante CM, Mondelli V. Childhood trauma and adulthood inflammation: a meta-analysis of peripheral C-reactive protein, interleukin-6 and tumour necrosis factor-alpha. Mol Psychiatry. 2016;21(5):642–9.

27. Lu S, Peng H, Wang L, Vasish S, Zhang Y, Gao W, et al. Elevated specific peripheral cytokines found in major depressive disorder patients with childhood trauma exposure: a cytokine antibody array analysis. Compr Psychiatry. 2013;54(7):953–61.

